# Hydrogel-embedded precision-cut lung slices support *ex vivo* culture of *in vivo*-induced premalignant lung lesions

**DOI:** 10.1101/2024.04.29.591698

**Authors:** Caroline Hauer, Rachel Blomberg, Kayla Sompel, Chelsea M. Magin, Meredith A. Tennis

## Abstract

Lung cancer is the leading cause of global cancer death and prevention strategies are key to reducing mortality. Medical prevention may have a larger impact than treatment on mortality by targeting high-risk populations and reducing their lung cancer risk. Premalignant lesions (PMLs) that can be intercepted by prevention agents are difficult to study in humans but easily accessible in murine preclinical carcinogenesis studies. Precision-cut lung slices (PCLS) are underutilized as an *ex vivo* model for lung cancer studies due to limited culture time. Embedding PCLS within bioengineered hydrogels extends PCLS viability and functionality for up to six weeks. Here, we embedded PCLS generated from urethane-induced murine PMLs in cell-degradable and non-degradable hydrogels to study viability and activity of the tissues over six weeks. PMLs in hydrogel-embedded PCLS maintained viability, gene expression, and proliferation. Treatment of hydrogel-embedded PCLS containing urethane-induced PMLs with iloprost, a known lung cancer prevention agent, recapitulated *in vivo* gene expression and activity. These studies also showed that iloprost reduced proliferation and PML size in hydrogel-embedded PCLS, with some differences based on hydrogel formulation and suggested that hydrogel-embedded PCLS models may support long-term culture of *in vivo* generated PMLs to improve preclinical studies of lung cancer and prevention agents.

## Introduction

Lung cancer is still the leading cause of cancer death in the United States among men and women and, while promising new therapies have been developed in recent decades, prevention remains the best strategy for reducing mortality from lung tumors.^1^ Avoiding tobacco smoke exposure is the best behavioral prevention for many cancers, especially lung cancer, but in high-risk populations, such as former smokers or cancer survivors, medical prevention approaches could have a significant impact on risk reduction. Historically, lung cancer prevention trials based on large population data sets, such as β-carotene, were not successful, while trials based on preclinical evidence, such as iloprost, demonstrated effects on intermediate endpoints like premalignant lesion (PML) histology.^2, 3^ Accessing PMLs in human lung for mechanistic studies is challenging, reinforcing the critical role of preclinical studies in understanding the biology of PML progression and testing prevention agents.^4^ Mouse studies relevant to humans, such as those incorporating tobacco carcinogens, provide access to multiple time points along the progression spectrum, however, these can be lengthy and costly.^5, 6^ Improvements in preclinical modeling are necessary to enhance research efficiency and to speed clinical translation of lung cancer prevention agents.

Precision cut lung slice (PCLS) models are used often to study a variety of lung pathologies and treatments, including fibrosis, infections, and chronic obstructive pulmonary disease (COPD).^7-9^ *Ex vivo* PCLS studies for lung cancer have been limited to a few publications investigating lung tumor biology and therapy, as well as a recent study from our group exploring PCLS as a model of lung premalignancy.^10-15^ PCLS are generated from lung tissue, most commonly mouse or human, by infusing tissue with agarose, slicing it, and culturing the slices for experiments and analysis.^16^ Other 3D models use single cell types or co-cultures, but do not include multiple native cell types or maintain natural architecture.^17^ Cultured PCLS retain most cell types from the original tissue, have expected responses to external stimuli, and keep tissue structures intact. An added benefit of PCLS for animal studies is the ability to screen interventions *ex vivo* for efficacy prior to initiating *in vivo* studies, reducing animal number and burden. The primary challenge of using PCLS for lung cancer studies is that longevity of cultures was limited to one to two weeks, which did not allow time for studies on initiation, progression, or regression.

Recently, the Magin research group published an innovative approach to support PCLS *ex vivo* using tissue-engineering strategies.^18^ Lung tissue slices were embedded within engineered hydrogel biomaterials designed to extend maintenance of tissue architecture and viability for up to three weeks *ex vivo*.^18^ In a subsequent study, we maintained the health of carcinogen-exposed lung tissue slices for an unprecedented six weeks in culture with hydrogel embedding and promoted cellular proliferation and altered gene expression.^13^ Here, our collaborative group presents a study investigating the capacity of hydrogel-embedded PCLS to maintain lung PMLs generated by an *in vivo* urethane adenocarcinoma carcinogenesis model. In the present study, the non-degradable hydrogel that best supported initiation of premalignant cellular responses was compared to a hydrogel degradable by matrix metalloproteinase-9 (MMP-9). MMP-9 is an enzyme secreted by pulmonary cells and immune cells that is active during embryonic development for lung branching morphogenesis and angiogenesis, processes that may be aberrantly activated during lung cancer, and is associated with all stages and poor prognosis of lung cancer.^19-21^ Proliferation and gene expression within PMLs induced *in vivo* were maintained *ex vivo* and differences were measured between non-degradable and degradable embedding hydrogels. Introduction of the prevention agent iloprost to hydrogel-embedded PCLS changed gene expression, decreased proliferation, and reduced lesion area to different degrees non-degradable versus degradable hydrogels. Our results demonstrate a new model for studying the biology and interception of lung PMLs and how these processes are influenced by hydrogels that mimic the tissue microenvironment.

## Materials and Methods

*Generation of mouse lung premalignant lesions*: Wild type female A/J mice (eight weeks old, Jackson Laboratories) were housed in a pathogen-free facility in the University of Colorado Anschutz Medical Campus Vivarium. At 10 weeks old, mice were injected IP with 100 μl of 1 mg urethane/gram body weight dissolved in 0.9% saline vehicle or 100 μl 0.9% saline vehicle to induce PMLs associated with development of adenocarcinoma.^5^ Mice were weighed daily for seven days after urethane injection and weekly for the remainder of the experiment, per standard protocols, and no unexpected weights changes were observed. 13 or 15 weeks after urethane exposure, mice were euthanized with carbon dioxide and exsanguination and lungs were harvested for PCLS. Animal numbers for urethane exposure were based on expected lung slice/punch yield and punch number was determined by endpoint assay needs. All animal experiments and procedures were reviewed and approved by the University of Colorado Anschutz Medical Campus IACUC (Protocol #1085).

### PCLS generation

Lungs were cleared with a cardiac perfusion of 10 ml sterile phosphate-buffered saline (PBS; Gibco) through the right ventricle. Immediately after, the lungs were filled via tracheal perfusion with 1 ml of 1.5% low melting point agarose (Fisher Scientific) dissolved in sterile 4-(2-hydroxyethyl)-1-piperazineethanesulfonic acid (HEPES) buffer. The whole mouse was placed on ice for 10 min before the lungs were extracted and placed in 1 ml of DMEM media with 0.1% Penicillin/Streptomycin/Fungizone (P/S/F) antibiotics/antimycotics. Lung lobes were then cut into 500 mm slices with a vibratome (Campden Instruments) at 12 mm/s speed and standardized punches were created with a biopsy punch (4 mm, Fisher Scientific). The punches were washed with media three times, with 30-minute incubations at 37°C between each wash to remove the agarose. Punches were cultured overnight in 24-well plates before hydrogel embedding.

### PEG-Norbornene (PEG-NB) synthesis

Norbornene-conjugated eight-arm 10 kg/mol PEG was generated as described previously.^22, 23,3^ The purified product was characterized by 1H NMR (Bruker DPX-400 FT NMR spectrometer, 300 MHz) and only product that was at least 90% functionalized was used in experiments.

### Hydrogel formulations and rheology

Non-degradable hydrogel solutions consisted of PEG-NB macromer (7 wt%), dithiothreitol (DTT) crosslinker (thiol:ene = 0.9; Fisher), and the cellular adhesion peptides mimicking binding sites on fibronectin (CGRGDS; 0.1 mM), laminin (CGYIGSR; 0.2 mM), and collagen (CGFOGER; 0.1 mM) from GL Biochem. Degradable hydrogel solutions consisted of PEG-NB (7.5 wt%), an MMP-9 degradable peptide crosslinker (Ac-GCRD-VPLSLYSG-DRCG-NH2; thiol:ene = 0.9; GL Biochem), and the same cellular adhesion peptides.^24^ For photopolymerization, 2.2 mM of the photoinitiator lithium phenyl-2,4,6-trimethylbenzoylphosphinate (LAP; Sigma Aldrich) was added. The elastic modulus of both formulations was determined by parallel plate rheology as previously described.^25, 26^

### Hydrogel embedding and PCLS culture

An 8-mm diameter silicone mold facilitated hydrogel embedding as previously described.^13, 27^ First, 25 μl of the hydrogel precursor solution was added to the silicone mold, and polymerized by exposure to 365 nm UV light at 10 mW/cm^2^ for 5 min. Hydrogel formulations were designed to be slightly off-stoichiometry (thiol:ene = 0.9) to leave free reactive groups within this base layer after polymerization that enables the second hydrogel layer to covalently bond to the first. A 4-mm PCLS punch was placed on first hydrogel layer and covered with an additional 25 μl of hydrogel solution. The entire construct was exposed to 365 nm UV light at 10 mW/cm^2^ for 5 min to complete the final polymerization step, resulting in the PCLS punch being fully encapsulated in hydrogel on all sides. Hydrogel-embedded PCLS were gently transferred to culture media in 24-well plates and maintained at 37°C with 5.0% CO_2_ in DMEM:F12 (1:1; Gibco) media with 0.1% fetal bovine serum (Gibco) and 0.1% 100X P/S/F antibiotics. Media was changed every 48 hours. In chemoprevention experiments, hydrogel-embedded PCLS cultures were treated with 10 μM iloprost (Cayman Chemical) or 9 mM methyl acetate vehicle control (Sigma Aldrich) every 48 hours.

### Presto blue metabolic activity assay

Hydrogel-embedded PCLS were incubated in 500 μl of Presto blue reagent (Fisher Scientific; 1:10 dilution in culture media) for 2 hours at 37°C. Following incubation, 150 μl samples of presto blue solution were transferred, in triplicate, to a clear 96-well plate and fluorescence intensity measured at 520 nm excitation (Glomax plate reader; Promega). Fresh Presto blue solution was also read as a blank control, and average blank control values for each plate were subtracted from individual fluorescence intensity values. Weekly measurements were normalized to the initial reading.

### EdU incorporation assay

A 5-ethynyl-2′-deoxyuridine (EdU) incorporation assay was performed according to the manufacturer’s protocol (Invitrogen). Hydrogel-embedded PCLS were incubated for 16 hours in complete culture media with 10 μM EdU. Samples were then fixed (4% PFA), permeabilized (0.5% Triton X-100), and incubated in Click-iT reaction cocktail containing AlexaFluorazide for 30 min. Nuclei were then counterstained with 5 μM Hoechst 33342. Samples were imaged on an upright epifluorescent microscope (Olympus BX63) by acquiring a z-stack at 10x magnification and then performing Weiner deconvolution. In Fiji (ImageJ), single channel z-stacks were used to generate a maximum projection, thresholded to exclude background, and number of nuclei determined by counting particles. The number of EdU+ nuclei was divided by the number of total (Hoechst+) nuclei to calculate percent proliferating cells.

### RT-qPCR

Three to five hydrogel-embedded PCLS of the same experimental group were pooled together to form each sample, and RNA was extracted with the RNeasy Plus kit (Qiagen). qPCR was conducted on a CFX96 Touch (Bio-Rad) using qPCR Prime PCR Assays (Bio-Rad) (*ces, e-cadherin, cox2, il6*, and *vimentin*) and the standard protocol for SsoAdvanced SYBR Green Master Mix (Bio-Rad). PCR was conducted in triplicate however, some samples did not amplify in specific assays, as represented by dots in bar graphs. All gene expression data were normalized to the reference gene *rps18* and fold changes were calculated using the 2-ΔΔCt method.

### Cryosectioning

Hydrogel-embedded PCLS were fixed (4% paraformaldehyde; (Electron Microscopy Sciences) and quenched (100 mM glycine; Fisher) for both at room temperature with rocking for 30 min. Samples were washed three times with PBS, excess hydrogel was trimmed from around the tissue, and then were placed in optimal cooling temperature (OCT) compound to perfuse at room temperature for 48 hours. OCT-infused samples were transferred to 10×10×5 mm cryomolds (Sakura), with 3-5 samples stacked into one mold, before being flash frozen by submersion in liquid nitrogen-cooled 2-methylbutane (Sigma-Aldrich). Blocks were stored at -80°C until cryosectioning on a Leica CM1850 cryostat. Frozen blocks were removed from the mold and mounted with one side pressed to the specimen disc, such that each cryosection would contain parallel cross-sections of the stacked PCLS punches. Serial sections of 10 μm thickness were acquired at –20°C and stored at -80°C for further processing.

### Immunofluorescence staining

Frozen sections were thawed to room temperature and equilibrated for 3 min in PBS before being circled by an ImmEdge hydrophobic pen (Vector Laboratories). Sections were blocked for 45 minutes in 5% BSA in PBS and then incubated with primary antibodies in 5% BSA overnight at 4°C. Primary antibodies were rabbit anti-TTF1 (Abcam, ab227652) and chicken anti-KRT5 (BioLegend, 905901). Slides were washed three times (in 0.1% Tween-20 in PBS (PBST) before incubation with secondary antibodies (5 μg/ml; Invitrogen) in 5% BSA in PBS for 1 h at room temperature. After another three washes in PBST, sections were incubated in 300 mM DAPI for 15 min at room temperature and then washed a final time in PBS. Sections were mounted under a 24 x 50 mm coverslip in ProLong Gold antifade reagent (Invitrogen) and allowed to cure overnight before imaging.

### Peroxisome proliferator activated receptor gamma (PPARγ) activity assay

PPARγ activity was assessed using a PPARγ response element (PPRE) luciferase assay as described previously.^10, 13^ Immortalized Human Bronchial Epithelial Cells (HBEC; a gift from John Minna) were transfected using TransIT-X2 reagent (Mirus Bio). Conditions included 45 ng PPRE luciferase (a gift from Bruce Spiegelman; Addgene plasmid #1015) and 5 ng renilla control reporter vector (Promega), mock luciferase, and empty vector. 24 hours after transfection, 15 μL of PCLS-conditioned media was collected from iloprost or vehicle-treated PCLS and was added to HBEC cells. HBEC cells were incubated for an additional 24 hours and then luciferase activity was measured using the Dual-Luciferase Reporter assay kit (Promega) on a Glomax instrument (Promega). PPRE firefly activity was normalized to renilla activity and analyzed relative to vehicle controls.

### Live cell imaging

For weekly imaging of live hydrogel-embedded PCLS, samples were cultured in glass-bottom 24-well plates (MatTek Life Sciences). PCLS were stained in 250 μl of 5 μg/ml Hoechst 33342 and 2.5 μg/ml AlexFluor 647-conjugated anti-EpCAM (Biolegend 118212) by incubating for 1 hr at 37°C. Samples were washed 3x in PBS and then allowed to rest before imaging for 1 hr in 500 μl complete culture media. Immediately before imaging, 400 μl of media was removed, ensuring that the hydrogel-embedded PCLS were resting on the glass bottom of the well. Imaging was performed on an inverted fluorescent microscope (Keyence) at 4x magnification, and then fresh media was added back to each well and samples returned to long-term culture. Images of the same lesions were captured weekly, and lesion size was quantified in Fiji (ImageJ) by converting each dual-channel image to 16-bit grayscale, thresholding the positive signal, drawing a circular region of interest (ROI) around the lesion to exclude high signal coming from normal airways, and then finding the pixel area of the thresholded region. Data are presented a percent change of each lesion relative to its initial starting size.

### Statistical analysis

Experiments with two groups and a single independent variable were assessed by two-tailed student’s t-test. Dose responses with three or more groups and a single independent variable were assessed by one-way ANOVA with a Tukey’s test for multiple comparisons. Experiments with two independent variables were assessed by two-way ANOVA with a Tukey’s test for multiple comparisons. All statistical analyses were conducted in GraphPad Prism software (GraphPad Software, Inc., San Diego, CA)

### Data Availability

The data generated in this study are publicly available at

## Results

### Hydrogel embedding supports long-term culture of hydrogel-embedded PCLS with in vivo induced PMLs

Lung tissue containing PMLs and vehicle controls were generated by injecting A/J mice with urethane or saline, respectively, and harvesting lung tissue at between 13 to 15 weeks (Fig. 1A). Urethane is a tobacco carcinogen that consistently induces a KRAS mutation in mice, followed by initiation of PMLs and progression to carcinoma.^5^ All PCLS from urethane-exposed lung tissue contained a piece of a PML. After slicing, PCLS were embedded in non-degradable or degradable PEG-NB hydrogel formulations containing peptide sequences mimicking collagen, laminin, and fibronectin (Fig. 1B) and cultured at 5% CO_2_ and 37°C for six weeks. Rheology results showed the elastic modulus (stiffness) of non-degradable and degradable hydrogels were not statistically different and matched the stiffness previously determined to facilitate *ex vivo* induction of PMLs (Fig. 1C).^13^ Viability measurements were collected weekly using a Presto Blue assay. Embedding PML-containing PCLS in either non-degradable or degradable hydrogels supported viability of both urethane-exposed tissue with PMLs and saline-exposed control tissue through six weeks (Fig. 2A).

**Figure 1.**
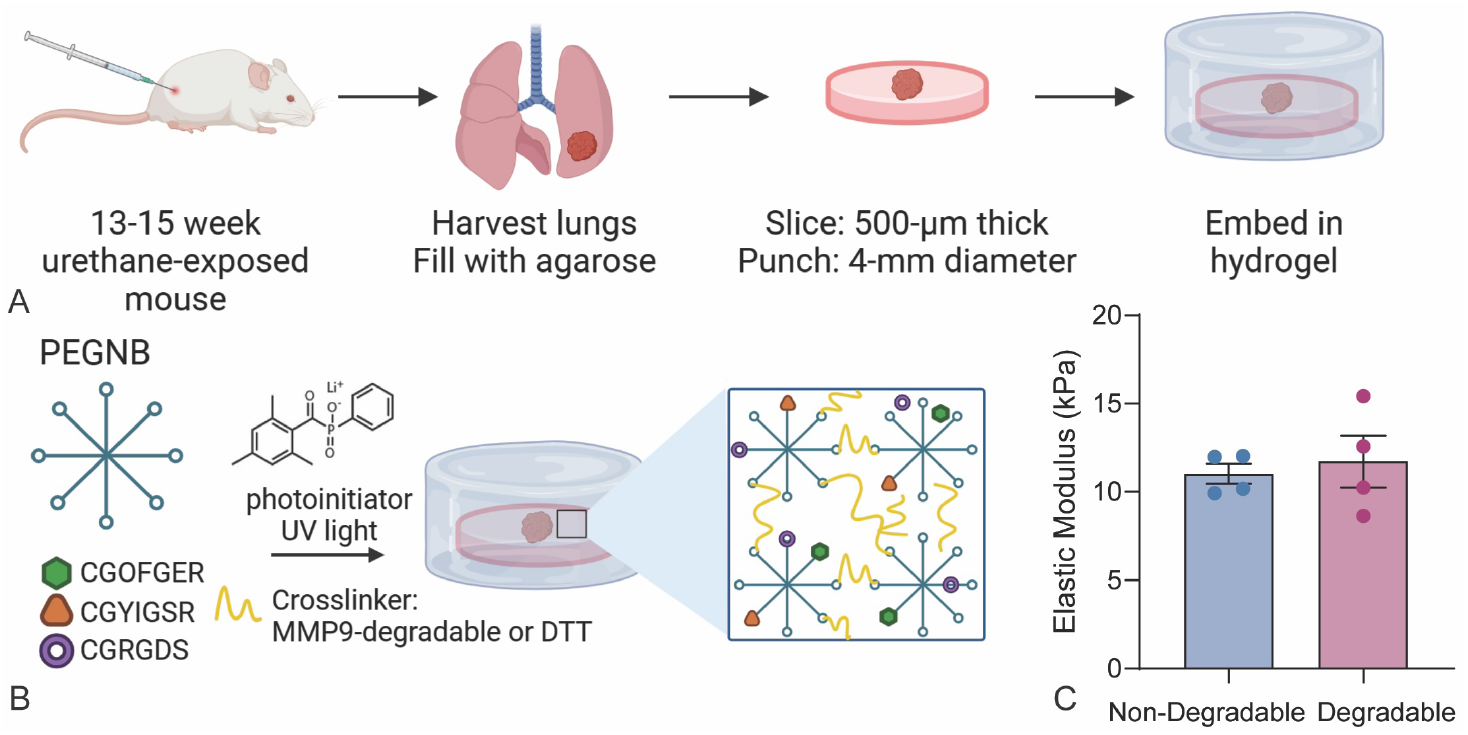
Generation of PML-containing PCLS, hydrogel composition, and rheology results. A) The process of PCLS generation from urethane-exposed mice included lung harvest, agarose filling of lungs, slicing filled lungs, and hydrogel embedding before culture. B) Two hydrogel compositions included PEG-NB, binding peptides (CGOFGER, CGYIGSR, CGRGDS), degradable or non-degradable) crosslinker, and a photoinitiator for photopolymerization. C) Rheology measurement of the elastic modulus (kPa) of non-degradable and degradable hydrogels (N=4; no statistical differences, student’s t-test). Error bars are SEM.

**Figure 2.**
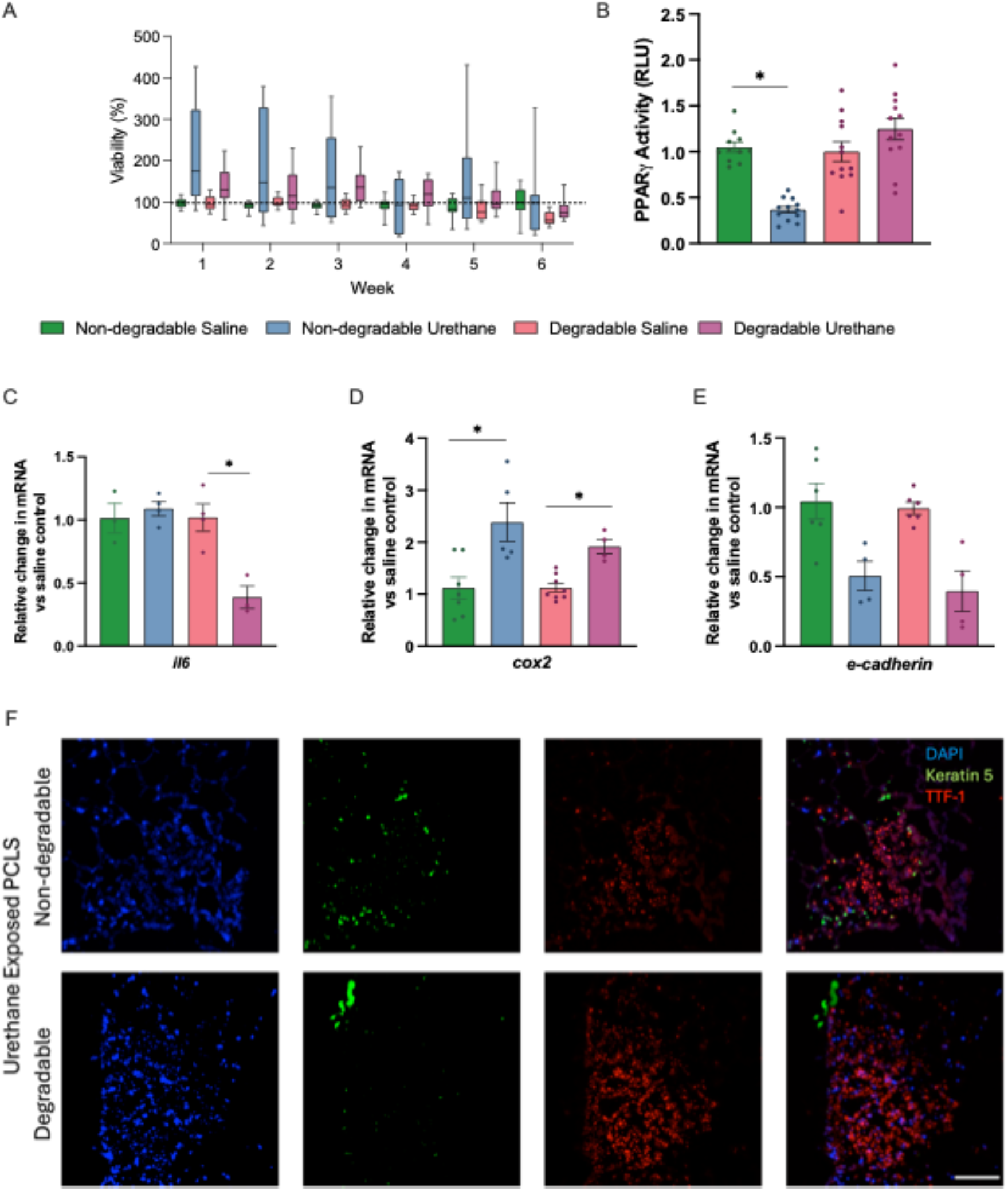
*In vivo* induced PMLs are viable and maintain expected characteristics in hydrogel-embedded PCLS. PCLS from *in vivo* urethane- or saline-exposed lung tissue were embedded in non-degradable or degradable hydrogel and cultured for six weeks. A) Weekly viability over six weeks, measured by Presto Blue assay. B) PPARγ activity in PCLS media at six weeks, measured by PPRE luciferase assay. C, D, E) qPCR measurement of gene expression for C) *il6*, D) *cox2*, and E) *e-cadherin* in PCLS at six weeks. qPCR results are normalized to *rps18* and presented as relative to saline control. F) DAPI (blue), Keratin 5 (green), and TTF-1 (red) staining in PCLS at six weeks. Scale bar is 100μM. Error bars are SEM. Scale bar 100 μm. Significance was determined by Student’s t-test and * is a p-value ≤0.05.

To compare hydrogel-embedded PCLS containing PMLs to data from *in vivo* lung tissue containing PMLs, we measured gene expression and PPARγ activity using a PPRE luciferase assay. In serum from the urethane model, PPARγ activity is reduced by urethane and in lung lesions and tissue, expression of inflammatory genes, such as *cox2* and *il6*, are increased, while epithelial genes, such as *e-cadherin*, are reduced.^28-31^ We found that at six weeks of culture, PPARγ activity was reduced in hydrogel-embedded PCLS media from *in vivo* urethane-exposed lung tissue compared to in vivo saline-exposed lung tissue (Fig. 2B). There was no change in PPARγ activity in media from PCLS embedded in degradable hydrogels. Expression of *il6* did not change in urethane-exposed PCLS embedded in non-degradable hydrogel, but was decreased in PCLS embedded in degradable hydrogel (Fig. 2C). Expression of *cox2* was increased and *e-cadherin* was decreased from *in vivo* urethane-exposed PCLS compared to *in vivo* saline-exposed PCLS embedded in both types of hydrogel (Fig. 2 D,E). TTF-1, a transcription factor for lung-specific proteins, identifies proliferating lung adenocarcinoma. In PCLS from *in vivo* urethane-exposed lung tissue, we measured TTF-1 expression by immunofluorescence and found it is maintained at six weeks (Fig. 2F). These data collectively suggest that lung tissue with PMLs from urethane-exposed mice can be cultured for six weeks in hydrogel-embedded PCLS with maintenance of viability and activity.

### Lung tissue with PMLs in hydrogel-embedded PCLS recapitulates in vivo gene expression and signaling with chemoprevention

To compare the effects of a chemoprevention agent in hydrogel-embedded PCLS to data from *in vivo* treated lung tissue, we measured gene expression and PPARγ activity. The lung cancer chemoprevention agent iloprost has been effective in numerous preclinical studies in mice, as well as in a phase II clinical trial of prevention in former smokers.^3, 29, 32^ Iloprost is a prostacyclin analogue that acts in part through PPARγ but likely has additional unknown targets related to its prevention activity.^32, 33^ We generated PCLS from urethane-exposed mouse lung tissue, embedded PCLS in non-degradable or degradable hydrogel, and then treated *ex vivo* with 10μM iloprost or vehicle every 48 hours for six weeks. Each PCLS form urethane-exposed lungs contained a piece of a PML. Viability of the PCLS was maintained over six weeks of culture in non-degradable and degradable hydrogels. (Fig. 3A).

**Figure 3.**
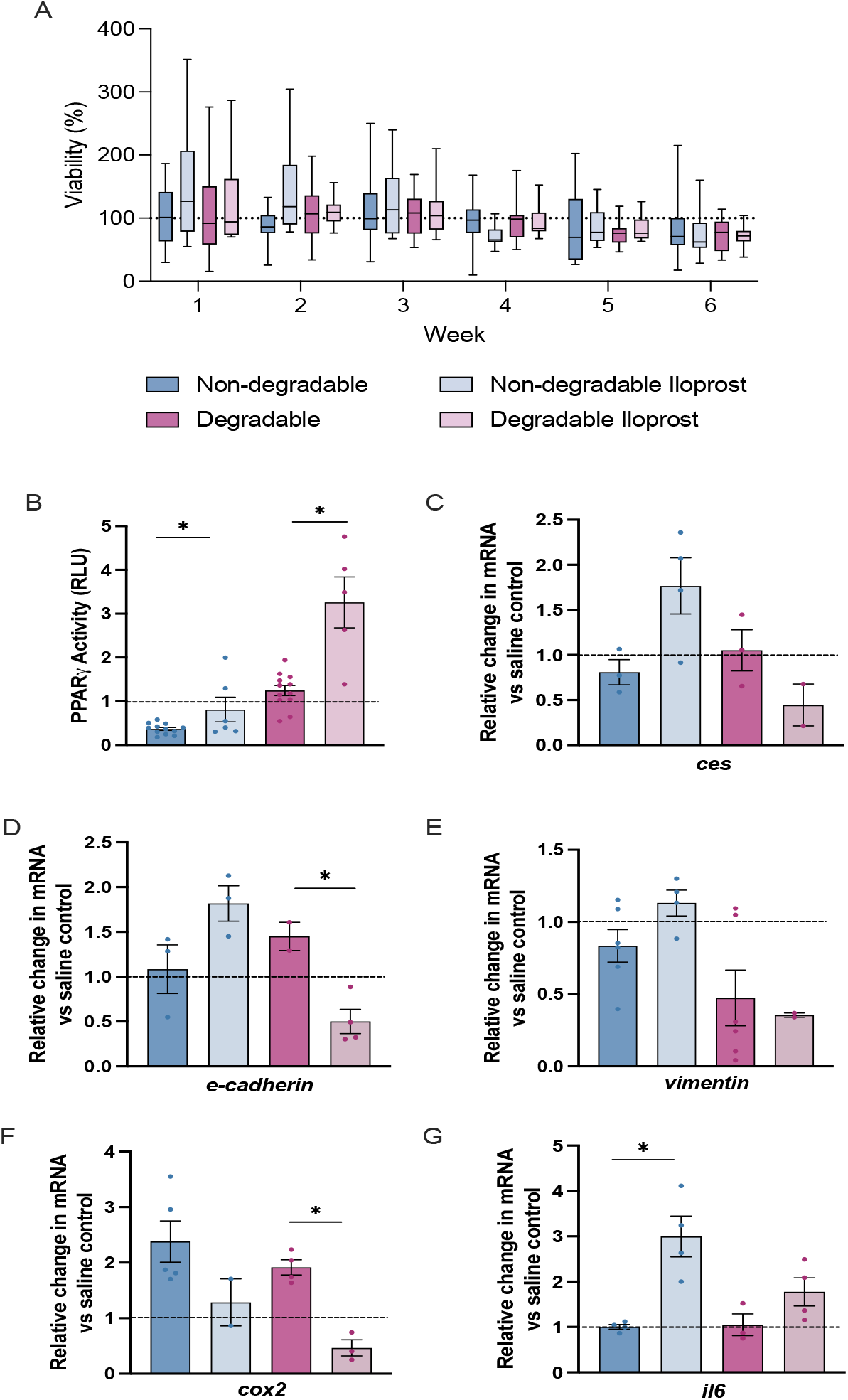
Hydrogel embedding supports response to ilo-prost chemoprevention in PCLS with urethane-induced PMLs. PCLS were generated from *in vivo* urethane-exposed lung tissue, embedded in non-degradable or degradable hydrogel, and treated with iloprost or control over six weeks of culture. A) Viability of PCLS, measured by Presto Blue assay. B) PPARγ activity in PCLS media, measured by PPRE luciferase assay. C-G) qPCR measured expression of C) *ces*, D) *e-cadherin*, E) *vimentin*, F) *cox2*, and G) *il6* and results are normalized to *rps18*. All results are presented relative to the hydrogel-specific saline vehicle controls (dotted line). Error bars are SEM. Significance was determined by one-way ANOVA and * is p-value ≤0.05.

Measurements of PPARγ activity in the hydrogel-embedded PCLS media revealed that in both types of hydrogel, PPARγ activity increased in iloprost-treated samples compared to vehicle control (Fig. 3B). Iloprost alters gene expression *in vivo* to mitigate effects of carcinogen exposure, including increasing *e-cadherin* and *carboxylesterase* (*ces*) and decreasing *vimentin* and *cox2*.^10, 30, 34^ Here, we found that iloprost increased *ces* compared to vehicle in non-degradable hydrogel PCLS (p=0.06) but not in degradable hydrogel PCLS (Fig. 3C). Iloprost increased *e-cadherin* expression in non-degradable hydrogel-embedded PCLS (p=0.09) but decreased it in degradable hydrogel PCLS (Fig. 3D). *Vimentin* expression did not significantly change in either hydrogel condition but *cox2* decreased in both with iloprost compared to vehicle control (Fig. 3 E,F). In studies of lung and heart disease, iloprost either decreases or does not affect IL-6; however, in short-term PCLS experiments, we previously found increased IL-6 with iloprost treatment.^10^ *Il6* expression increased with iloprost in both hydrogel types (Fig. 3G). Expected changes in gene expression and signaling with *ex vivo* iloprost treatment of *in vivo* urethane-exposed hydrogel-embedded PCLS were observed, suggesting that chemopreventive agents were active in hy-drogel-embedded PCLS and that hydrogel composition that influence the microenvironment may lead to different effects.

### Chemoprevention in hydrogel-embedded PCLS reduces the proliferation and size of PMLs

To demonstrate the efficacy of a chemoprevention agent on PMLs in hydrogel-embedded PCLS, we measured proliferation and size of PMLs during culture. Urethane-induced lesions *in vivo* have increased proliferation ^35, 36^ In *in vivo* studies, iloprost reduced PML number and in a human study, reduced PML grade and proliferation.^3, 29, 32^ We generated PCLS containing PMLs from urethane- or saline-exposed mice, embedded these samples in non-degradable or degradable hydrogel, and exposed the cultures to 10μM iloprost or vehicle control every 48 hours for six weeks. Analysis of proliferation by EdU at six weeks showed a trend toward decreased proliferation in iloprost-treated, hydrogel-embedded PCLS (Fig. 4A,B). In the degradable hydrogel at three weeks, proliferation was significantly higher in the PCLS from urethane-exposed lung tissue and significantly decreased with *ex vivo* iloprost treatment. At six weeks, proliferation was lower but still maintained a pattern of increased proliferation in hydrogel-embedded PCLS from urethane-exposed, PML-containing lung tissue and decreased in iloprost-treated, hydrogel-embedded PCLS.

**Figure 4.**
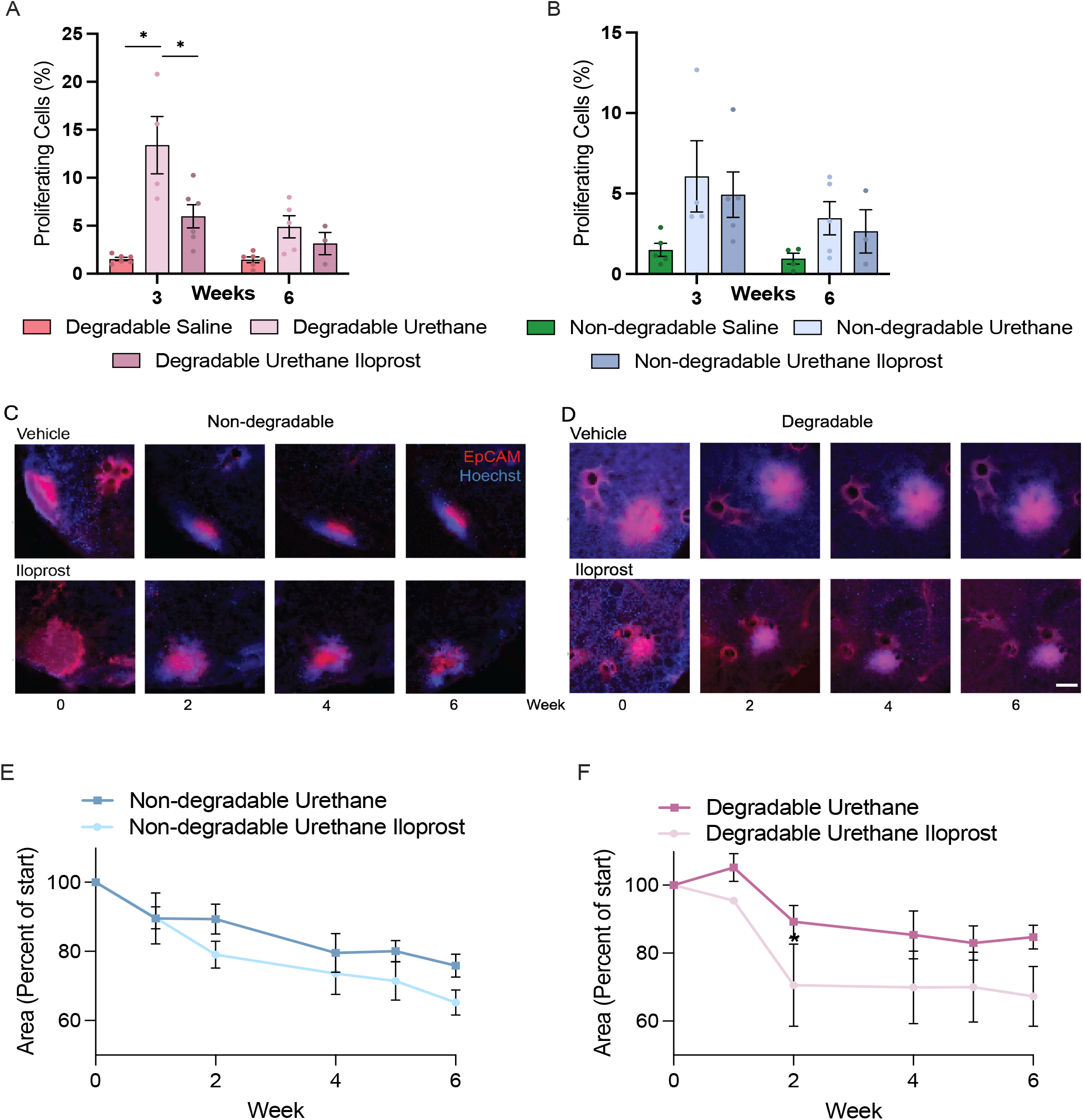
Iloprost treatment of urethane-induced PMLs in hydrogel-embedded PCLS reduces proliferation and size. PCLS were embedded in hydrogel and cultured for six weeks. A, B) Proliferation was measured by EdU and quantified as percent proliferating cells in PCLS embedded in A) non-degradable hydrogel or B) degradable hydrogel. C, D) PML size in PCLS was measured with live cell imaging using EpCAM and Hoescht staining every two weeks in PCLS embedded in C) non-degradable or D) degradable hydrogel. Scale bar is 100μm. E, F) Quantification of live cell imaging as percent area of PML starting area in PCLS embedded in E) non-degradable or F) degradable hydrogel. Significance was measured by two-way ANOVA and * is p≤0.05. All assays were conducted in triplicate and error bars are SEM.

Real-time live cell imaging with an anti-EpCAM antibody monitored the size of PMLs over six weeks with measurements at 1, 2, 4, 5, and 6 weeks (Fig. 4 C, D). In PCLS embedded in the non-degradable hydrogel, PML area significantly decreased with iloprost treatment at weeks 2, 4, 5, and 6 (p=0.01, 0.0006, 0.0002, and 0.0001, respectively) compared to time zero (Fig. 4E). In the urethane only samples, a significant decrease in size was only observed at weeks 5 and 6 (p=0.04 and 0.007, respectively) compared to time zero (Fig. 4E). The final difference in area between iloprost and control was about 10% (p=0.08). At week one, the area of PMLs in the degradable hydrogel, urethane only PCLS increased and, while they decreased in area after this timepoint, these PMLs remained the largest over the six weeks of culture (Fig. 4F). Significant differences in area compared to time zero were not observed in the urethane only PMLs in degradable hydrogel, while iloprost-treated PMLs had significant decreases in area at weeks 4, 5, and 6 (p= 0.05, 0.05, 0.03, respectively) compared to time zero. The area of iloprost treated PMLs in the degradable hydrogel decreased at a faster rate than the urethane only, with a difference of about 20% at six weeks (p=0.07). In the degradable hydrogel, PML area in both iloprost and urethane only PCLS decreased at a slower rate than in the non-degradable hydrogel. Overall, these results suggest that treatment with iloprost reduces proliferation and PML area in hydrogel-embedded PCLS and that degradable hydrogels better maintained characteristics of premalignancy over time.

## Discussion

The results presented here demonstrate that lung PMLs induced *in vivo* by urethane can be maintained in hydrogel-embedded PCLS for six weeks. Gene expression and signaling activity in hydrogel-embedded PCLS cultured for six weeks recapitulated previous results from *in vivo* lung models.^29, 30, 37^ *Ex vivo* prevention agent treatment of PMLs in hydrogel-embedded PCLS mimicked results from *in vivo* treatments, with changes in gene expression, signaling activity, proliferation, and PML size associated with treatment.^29, 30, 37^ Human lung cancer progression is associated with the deposition of dense extracellular matrix and increases in tissue stiffness, which supports epithelial and tumor cell proliferation.^38, 39, 40^ In agreement with this *in vivo* phenomenon, we previously found that a stiff hydrogel incorporating the collagen-derived peptide GFOGER best supported the initiation of carcinogenesis by vinyl carbamate (a metabolite of urethane) in mouse PCLS.^13^

In this study, we build on this work by using stiff hydrogels containing the GFOGER sequence, and interrogated whether changes in the hydrogel degradability might alter PML activity or growth. The DTT (synthetic) crosslinker used here is non-degradable, while the peptide (degradable) crosslinker can be acted upon by MMP-9 secreted by various cell types in PCLS. Increased MMP-9 expression is associated with all stages of lung cancer and poor prognosis, but also has been detected in mild squamous dysplasia, where cells expressing MMP-9 moved through disrupted basal membranes with colocalized collagen and MMP-9.^19, 41^ One goal of these experiments was to determine if the ability to degrade the hydrogel environment through MMP-9 activity would lead to effects on PMLs or response to treatment. In comparisons of the non-degradable and degradable hydrogels, differences in gene expression followed a similar pattern for urethane- and saline-exposed hydrogel-embedded PCLS in *e-cadherin* and *cox2*. In contrast, *il6* was significantly lower in urethane-exposed PCLS in degradable hydrogel but did not change in urethane-exposed PCLS in non-degradable hydrogel. In iloprost-treated, hydrogel-embedded PCLS, there were more differences in gene expression and PPARγ activity between hydrogel composition and some were not consistent with previous data. PPARγ activity was higher with iloprost in both hydrogel conditions compared to control, as expected, and was much higher in degradable hydrogel-embedded PCLS. Inhibition of MMP-9 occurs with PPARγ activation in lung tumor cell lines, a pathway that may be activated by increased availability of MMP-9-degradable material within degradable hydrogel-embedded PCLS conditions.^42-44^ Increased activity of PPARγ in MMP-9 degradable hydrogel-embedded PCLS supports our observation of a difference in proliferation and PML size with iloprost treatment in degradable hydrogels, as PPARγ activity is critical for effects of iloprost on lung tumor cells.^32, 33^ In contrast to the PPARγ data, *e-cadherin* and *ces* expression decreased with iloprost in degradable hydrogel-embedded PCLS, which does not agree with previous data.^30^

Chemical carcinogenesis models of lung adenocarcinoma in mice employ a range of carcinogens, but all use large numbers of mice and tend to have long tumor latency.^5, 6^ Testing prevention agents in mouse lung adenocarcinoma carcinogen models require even more animals to observe limited effect sizes.^29, 32, 45-47^ Despite challenges, preclinical studies are critical for advancing precision prevention approaches.^4^ Our new hydrogel-embedded PCLS model could reduce the number of animals required for preclinical prevention agent studies by supporting high throughput *ex vivo* experiments for hypothesis refinement or agent screening prior to full *in vivo* studies. This model also allows investigations into stages and mechanisms of lung PML development that are not accessible in humans and offers more opportunities for manipulation than *in vivo* models.^11, 12, 48^ Previous lung cancer PCLS studies measured response to agents in tumor or normal tissue for periods of several hours to several days, but none studied premalignant lesions or used long-term culture.^14, 49-51^ While other three-dimensional modeling systems may be useful to answer specific questions related to lung cancer, they have important limitations for lung premalignancy and prevention research. For example, patient-derived tumor organoids may be an effective tool for identifying tumor-specific treatments but these cultures require a large number of initials cells, lack resident immune cells, and normal airway cells often overgrow the tumor cells.^52^ Microfluidic lung-on-a-chip models can recapitulate physiologic and mechanical functions that affect lung cancer development, but disrupts the native environment of tumor cells.^53, 54^ The complete tissue architecture and maintenance of many biologic processes in the hydrogel-embedded PCLS model is critical for studying tissue level mechanisms and with further development this model will support critical contributions to lung cancer research.

Our study generated PCLS from one *in vivo* time point; additional time points could be added in future studies to assess how progressive stages can be supported by hydrogel-embedded PCLS. We previously demonstrated the presence of macrophages and T cells in hydrogel-embedded PCLS at six weeks.^10, 13^ Comprehensive characterization of the presence or activity of immune cells throughout the six week culture will support adoption of this model to investigate immune mechanisms and test immunoprevention agents. This study focused on adenomatous PMLs induced by urethane and we could apply this model also to squamous cell carcinoma (SCC) by using the NTCU mouse model to generate PCLS.^55, 56^ We have explored using human tissue in hydrogel-embedded PCLS for initiating PMLs *ex vivo*, but accessing human PMLs in tissue suitable for long-term PCLS is challenging.^13^ We continue to investigate the possibilities for human relevant hydrogel-embedded PCLS, including designing hydrogels to mimic human microenvironment conditions in mouse PCLS.

A hydrogel-embedded PCLS model that allows long term culture of lung PMLs is an exciting development in preclinical modeling of lung cancer that is expected to have a significant impact on PML biology and prevention agent studies. Future studies focused on bioengineering aspects could include modifications of hydrogel composition to mimic different parts of the PML microenvironment or altering hydrogel stiffness to parallel tissue stiffness in mouse of human lung cancer development. Future studies focused on cancer could investigate comorbidities with hydrogel-embedded PCLS from COPD or fibrosis mouse models that are exposed to lung cancer carcinogens *ex vivo*. Humans are often exposed to multiple carcinogens, but *in vivo* studies of combined carcinogens are challenging, so hydrogel-embedded PLCS could be used to explore biology with multiple exposures. While our focus has been on PMLs and prevention, this foundational PCLS model could also be used to study solid tumors and targeted therapies, broadening the potential impact of this model on lung cancer research.

## Author’s Contributions

C. Hauer: Investigation

R. Blomberg: Investigation, methodology, formal analysis, writing -original draft, writing-review & editing.

K. Sompel: Investigation

C.M. Magin: Conceptualization, funding acquisition, project administration, data curation, resources, supervision, and writing – review and editing.

M.A. Tennis: Conceptualization, funding acquisition, project administration, supervision, writing-original draft, writing-review & editing.

## Acknowledgements

This work was supported by funding from the University of Colorado Cancer Center Thoracic Oncology Research Initiative (to C.M.M and M.A.T.), the National Cancer Institute (R21CA252172 to R.B., C.M.M, and M.A.T.), and the NIH (5T3HL007085-47 to R.B.).

## Notes

Conflict of interest: C.M.M serves as the Vice Chair of the Board of Directors for the Colorado BioScience Institute. All other authors declare no potential conflicts of interest.

### Competing Interest Statement

C.M.M serves as the Vice Chair of the Board of Directors for the Colorado BioScience Institute.

